# BRG1 Cooperates with PLK1 to Regulate Dormant Origin Activation During S Phase

**DOI:** 10.1101/2025.09.27.678965

**Authors:** Saddam Hussain, Deepa Bisht, Rohini Muthuswami

## Abstract

The ATP-dependent chromatin remodeler BRG1 is well-established in the regulation of gene expression, DNA replication, homologous recombination, DNA damage repair, and apoptosis. While its genome-wide occupancy has primarily been linked to transcriptional regulation, its potential role in DNA replication control remains less well defined. In this study, we investigated BRG1’s involvement in replication origin regulation by integrating BRG1 ChIP-seq data with replication origin mapping and chromatin state analyses. Our results revealed that BRG1 is preferentially enriched at genomic regions characterized by DNase I hypersensitivity, low GC content, and a capacity to form G-quadruplex structures hallmarks of dormant replication origins. Notably, BRG1-bound sites were also enriched in late-replicating, heterochromatic regions. These findings suggest a specific association of BRG1 with dormant replication origins. To validate this association, we performed functional assays in HeLa cells. BRG1 depletion via siRNA led to reduced replication fork progression and increased activation of dormant origins, consistent with replication stress. Mechanistically, we found that BRG1 physically interacts with the mitotic kinase PLK1. Loss of BRG1 reduced chromatin-bound levels of both BRG1 and PLK1, implicating BRG1 in PLK1 recruitment or stability. Importantly, overexpression of PLK1 in BRG1-depleted cells rescued replication fork progression, supporting a cooperative role for BRG1 and PLK1 in regulating replication through dormant origins. Together, these data uncover a previously unrecognized function for BRG1 in the selective regulation of dormant replication origin firing, acting in concert with PLK1.

## INTRODUCTION

In humans, approximately 25,000–50,000 active replication origins are sufficient to duplicate the entire genome during a single cell cycle [1]. However, the number of potential replication origins in the genome far exceeds this requirement. High-throughput sequencing approaches—including Repli-seq, SNS-seq, ChIP-seq, and replication initiation sequencing—have identified an unexpectedly high number of replication origins across various human cell lines, with estimates reaching up to ∼320,748 [2][3] [4] [5] [6] [7]. A substantial proportion of these origins are not activated in every cell cycle. Instead, they are passively replicated by adjacent replicons and are referred to as dormant origins [8].

Despite their shared protein composition with active (or firing) origins, the mechanisms that govern the activation of dormant origins remain poorly understood. One key trigger is replication stress, which can arise from both endogenous and exogenous sources [9, 10]. Endogenous sources of replication stress include spontaneous DNA damage [11], repetitive DNA sequences [12], G-quadruplex structures [13], and replication-transcription collisions [14], whereas exogenous sources include genotoxic agents such as carcinogens and chemotherapeutic drugs. Under replication stress, replication forks may slow down or stall [15–17], leading to the activation of nearby dormant origins to ensure complete and timely genome duplication [9, 18].

Importantly, emerging evidence suggests that the choice between dormant and firing origins is influenced by the local chromatin environment and structural DNA features. Therefore, differentiating between firing and dormant origins involves analysis of specific genomic and epigenetic features [7, 19, 20]. Firing origins typically initiate replication early in S-phase and are enriched in regions of open chromatin, often upstream of gene promoters. These regions are characterized by active chromatin marks such as H3K4me2, H3K4me3, H3K9ac, H3K27ac, H2A.Z, and H4K20me1 [21–23][19, 24, 25]. In contrast, late-firing and dormant origins are more frequently associated with repressive histone modifications, including H3K9me3 and H3K27me3— the latter being a hallmark of Polycomb-repressive complex localization [23, 26, 27]. Additionally, origin density positively correlates with GC content [2, 19, 28], and regions rich in GC nucleotides tend to form G-quadruplex structures, especially in nucleosome-free zones, influencing both replication and gene regulation [29]. A recent study demonstrated that most folded G-quadruplexes co-localize with BRG1-occupied genomic sites [30].

The chromatin environment surrounding replication origin is critical not only for origin activation but also for preventing re-replication within the same cell cycle. Chromatin remodelers dynamically regulate this environment during key cellular processes such as replication [31], transcription [32, 33], and DNA repair [34, 35]. Among these, the SWI/SNF ATP-dependent chromatin remodeling complexes have been implicated in origin activation in both *S. cerevisiae* and mammalian systems [36, 37]. BRG1 (SMARCA4), the ATPase subunit of the polybromo-associated BAF (pBAF) complex, contains an N-terminal ATPase domain essential for remodeling activity and a C-terminal bromodomain that binds acetylated lysines on histones to modulate chromatin structure [38, 39]. Beyond its role in transcription and DNA damage response, BRG1 has been shown to associate with replication assembly components such as ORC1 and the GINS complex [37], as well as Geminin, RPA1, and RPA2 [40], indicating a role in pre-replication complex (pre-RC) licensing and replication elongation. BRG1 also interacts with TOPBP1, a critical factor at replication forks during stress responses [41]. Functional studies demonstrate that BRG1 deletion downregulates the expression of ORC1 and CDC6 [42], resulting in G1 phase arrest [43], and alters replication timing domains in embryonic stem cells [17]. BRG1 loss induces replication stress, slows replication fork progression, and increases activation of dormant origins [44]. Conversely, BRG1 reconstitution restores fork velocity and reduces dormant origin firing [42, 45]. Notably, BRG1 mutations reduce fork speed by approximately 50% and its depletion decreases inter-origin distances by ∼35% [37], a phenotype comparable to depletion of replication stress regulators such as Timeless, Claspin, and Chk1 [46]

BRG1 forms a regulatory loop with another chromatin remodeler, SMARCAL1, during DNA damage response [47]. DNA double-strand breaks increase the expression of both BRG1 and ATR, suggesting that BRG1 is upregulated in response to genotoxic stress [48, 49]. BRG1 deficiency results in pan-nuclear γ-H2AX foci formation and triggers replication stress signaling via ATR and Chk1 activation [50, 51]. Similarly, PLK1, a serine/threonine kinase essential for cell cycle progression, plays a critical role under replication stress [52]. PLK1 forms a complex with replication machinery components including MCM2/7, HBO1, and ORC2 [53, 54], and its phosphorylation of HBO1 is necessary for pre-RC assembly and origin licensing [55]. Inhibition of PLK1 using BI2536, a known pharmacological inhibitor, delays S-phase progression (notably at 4– 6 hours post-G1/S release) by compromising ORC2 phosphorylation at Ser188 [54]. In unperturbed cells, PLK1 co-localizes with ORC1 and MCM2 on chromatin. PLK1 depletion impairs DNA replication during early and mid-S phase, while overexpression partially restores replication, particularly in mid to late S phase. Importantly, PLK1 overexpression does not elevate DNA damage markers such as γ-H2AX, but enhances replication rate by 1.3–1.5-fold in the absence of exogenous stress [56]. PLK1 is also implicated in the regulation of replication origin firing via its control over MTBP, TopBP1, and Rif1 expression, and its activity is required for the release of MTBP from chromatin during elongation [57].

To date, no single molecular marker uniquely defines dormant origins; instead, multiple epigenetic and genomic features are employed to map their genomic locations. While PLK1 depletion has been consistently associated with reduced replication progression and increased origin firing (suggesting activation of dormant origins), the role of BRG1 in this context has been explored less extensively, and no prior studies have addressed potential functional crosstalk between BRG1 and PLK1. The established role of BRG1 in transcription and DNA repair, combined with replication defects upon its depletion resembling those observed with PLK1 loss, hints at a regulatory interaction during S phase [42, 56–58].

Given that both BRG1 and PLK1 engage common replication factors, modulate ATR-Chk1 signaling, and are individually essential for normal replication fork progression and dormant origin regulation, we hypothesized that BRG1 might act as a chromatin-associated regulator of dormant origins through interaction with PLK1 during S phase. Using publicly available replication origin datasets alongside BRG1 ChIP-seq data, we show that BRG1 is enriched at dormant replication origins. BRG1 depletion via siRNA leads to impaired replication dynamics and activation of dormant origins, while PLK1 overexpression in BRG1-deficient cells rescues replication defects. These findings suggest that BRG1 serves as a chromatin-bound determinant of dormant origin regulation, acting through functional interplay with PLK1.

## MATERIAL AND METHODS

### Chemicals

All general laboratory chemicals, unless otherwise mentioned, were purchased from Thermo Fisher, USA, and Sisco Research Laboratories, India. Cell culture chemicals and reagents like DMEM, Trypsin-EDTA solution, Penicillin-Streptomycin-Amphotericin B (PSA) solution, and Bradford reagent were purchased from Hi-Media, USA. Thymidine, CldU and IdU were purchased from Sigma-Aldrich, USA. Restriction endonucleases, reverse transcriptase, and RNase inhibitor were purchased from New England Biolabs, USA. SYBR Green PCR Master mix was purchased from Kapa Biosystems USA. QIAquick gel extraction kit was purchased from Qiagen, USA. Protein-G fast flow bead resin, Immobilon-P PVDF membrane, luminol, and p-coumaric acid were purchased from Merck-Millipore USA. X-ray films, developer, and fixer were purchased from Kodak USA. TRIzol^®^ reagent and QIAprep^®^ Spin Miniprep Kit was purchased from Qiagen, Netherlands. Micro-amp Fast 96-well reaction plates, 0.1ml was purchased from Applied Biosystems USA. Cell culture grade dishes and serological pipettes were purchased from SPL Life Sciences, USA.

### Antibodies

BRG1 (Catalog# B8184, Abcam), PLK1 (ab17056, Merk), CldU (ab6326, Abcam), IdU (BD347580, BD biosciences), FITC-conjugated anti-rat IgG (Invitrogen 31629), TRITC-conjugated anti-Mouse IgG (Invitrogen A16071) and β-actin (Catalog# A1978) antibodies were purchased from Sigma-Aldrich (USA).

### Cell culture maintenance and treatments

HeLa cells were purchased from NCCS Pune, India. The cells were cultured in DMEM media in a humidified chamber at 37°C in the presence of 5% CO_2_. For all the experiments, HeLa cells were used after the third passage.

### siRNA and plasmid transfection

siRNA pool for *BRG1* and non-targeting control (NTC) was purchased from Dharmacon, USA. *BRG1* and full-length *PLK1*(FLPLK1) were cloned into pcDNA3.1 Zeo-LAP vector. For transfection, plasmids were isolated using a Qiagen plasmid purification kit, and the concentration was quantified using NanoDrop™ 2000 spectrophotometers. Transfection was done using 3 µl of si*BRG1*, siNTC and lipofectamine 3000 reagent as per manufacturer’s protocol. In experiments where siRNA was co-transfected with plasmids, 4 µg of respective plasmid DNA was used. The transfection mix was then added to the culture dish, and cells were grown for the indicated time points.

### Cell synchronisation and transfection

Briefly, HeLa cells were seeded to a confluence of 30-40% and incubated at 37°C for 12-16 hours. After cell transfection, thymidine was added to a final concentration of 2 mM only after 6 hours, and cells were left in the incubator for 18 hours. After 18 hours, the cell media was changed with fresh serum-containing media (SCM) for 8-9 hours. Cells were given a second thymidine block by adding 2 mM thymidine for 16 hours. After 16 hours, 0 hour samples were collected, and the rest of the cells were grown in fresh media for 4 more hours.

### Co-immunoprecipitation

HeLa cells were grown in a 100 mm culture grade dish to confluency of 70-80%. The cells were lysed in RIPA buffer for 15-20 min followed by sonication in a water bath. Collected supernatant was incubated with equilibrated protein AG beads (slurry) at 4°C for 1 hour. The supernatant was obtained by centrifuging the beads at 3000 rpm for 5 min, and protein was quantified using a Bradford reagent. ∼ 700 µg pre-cleared protein extract was added with antibodies and incubated at 4°C for 16 hours in a rotating wheel. The protein AG beads were blocked with BSA (250 µl slurry blocked with 0.1 µg/µl BSA) and incubated overnight at 4°C in a rotating wheel. The next day, protein antibody complexes were incubated with 40 µl BSA-blocked beads for 3 hours at 4°C. The supernatant was separated by centrifuging at 2000 rpm for 5 min, and the beads were washed 4 times (2000 rpm for 2 min at 4°C) with RIPA buffer. Next, the beads were resuspended in RIPA buffer, and the desired volume of protein loading dye was added. The sample was boiled for 10 min at 95°C in a heating block. The protein sample was loaded onto the SDS gel and proceeded for Western blotting.

### Western blotting

HeLa cells were grown in a 100 mm culture grade dish to a 30-40% confluency and blocked G1/S boundary with double thymidine treatment. Cell lysates were prepared by incubating them with RIPA buffer, and protein was quantified using a Bradford reagent. 40 µg of protein is loaded in 10% SDS-PAGE. The protein bands were transferred onto PVDF membrane. The membrane was blocked in 5% skimmed milk prepared in 1X PBST (with 0.1% Tween) for 1-2 hours at room temperature. The blocked membrane was incubated with primary antibodies at 4°C for 16 hours. After washing the blots 3 times with PBST for 5 min each membrane is incubated with HRP-conjugated secondary antibodies at room temperature for 1-2 hours. The blots were then washed with 0.1% PBST (1X) 3 times for 5 min each. The blots were kept in PBS and then developed in the dark by following Enhanced chemi-luminescence method. The western blots were quantitated using Image J software after developing.

### Pre-extraction immunofluorescence

HeLa cells were grown to 40-50% confluency, blocked in G1/S through double thymidine block and transfected with si*BRG1* and/or *PLK1* plasmid. For IdU incorporation, 250 µM of IdU was added at 3 hours 30 min post double thymidine for 30 min. For pre-extraction of cytosolic proteins, the cells were permeabilized with ice-cold 0.5% Triton X-100 prepared in PBS for 5 min in dark at 4°C and cells were fixed with ice-cold methanol for 10 min in dark at 4°C. The cells were blocked with 2% BSA prepared in PBS for 2-3 hours at room temperature. The cells were incubated with primary antibody prepared in ice-cold 2% BSA in PBS and incubated at 37°C for 1 hour. The cells were washed 5 times with 0.2% PBST (0.2% Triton X-100 in PBS) for 5 min each. The cells were further incubated with respective secondary antibodies, incubated for 45 min at room temperature, and washed 5 times with 0.2% PBST for 5 min each. The coverslips were mounted onto the glass slides with 10 µl 70% glycerol. The edges were sealed and observed under a Nikon A1R HD confocal microscope (Eclipse *Ti-E*, Nikon), Nikon, Japan.

### RNA isolation and quantification

RNA was extracted from untreated and treated HeLa cells as explained in Radhakrishnan et al. [59]. Briefly, **c**omplementary DNA (cDNA) was prepared using 2 µg of total RNA and 1µl random hexamer. The volume was made up to 12 µl with DEPC-treated autoclaved water, and the reaction mix was incubated at 65°C for 5 min and then at 4°C. For each sample, 8 µl of master mix (2 µl of 10X reaction buffer, 0.2 µl of reverse transcriptase enzyme, 0.2 µl of RNase inhibitor, 0.5mM dNTPs and water) was added. The sample was then incubated for 10 min at 25°C followed by 60 min at 42°C. A final incubation of 10 min at 72°C was given to terminate the reaction, and then the sample was gradually brought at 4°C in the PCR cycler.

### Real-time quantitative PCR (qRT-PCR)

qRT-PCR was performed on Applied Biosystems 7500 Real-Time PCR system. The prepared cDNA was diluted by adding an equal volume of water and the reaction was set using 1µl of cDNA. Each reaction mix contained 5 µl of 2X KAPA SYBR FAST qRT-PCR Master Mix, 100 pmol forward and reverse primers, and water to make the total volume to 10 µl. The sample was then incubated at 95°C for 20 seconds followed by qRT-PCR reaction for a total 40 cycles (95°C for 3 s, 60°C for 30 sec) and finally, the melt curve was plotted at 95°C. The raw data was normalized and quantified using the ΔΔCt method. The data was plotted using Sigma plot version 12.0.

### DNA fibre assay

Briefly, cells were seeded in 60 mm dishes to a confluency of 30-40%. As mentioned earlier, the cells were synchronised using double thymidine block and released from G1/S phase for 3 hours 30-40 min. The cells were sequentially pulse-labeled with 25 mM CldU and 250 mM IdU for 20 and 30 min, respectively. The cells were then harvested and resuspended in 50 ml of chilled PBS. The cell suspensions (3 µl) were placed on glass slides (Superfrost) and allowed to air-dry. The cell suspension was then mixed with 8 ml of lysis buffer [0.5% SDS, 200 mM Tris-HCl (pH 7.4), 50 mM EDTA] and incubated for 2 min. The slides were inclined at an angle between 25-45° to spread the suspension. Once dried, the DNA spreads were fixed by incubating the slides in a 3:1 solution of methanol-acetic acid for 10 min, followed by denaturation using 2.5 N HCl for 80 min. After 3-5 rinses in 1X PBS, the slides were incubated with blocking buffer (2% BSA and 0.01% Tween-20 in PBS) for 15 min. This was followed by incubation with rat anti-BrdU (CldU) antibody for 90 min in a humidified chamber at room temperature. The slides were washed once with 0.1% Tween-20 and fixed in 4% formaldehyde solution for 15 min. After five PBS washes, the slides were incubated with Anti-rat FITC secondary antibody. After incubation for 1 hour at room temperature, the slides were washed twice with 0.1% Tween-20 and incubated overnight with mouse Anti-IdU antibody at 4°C. The slides were washed twice with PBS and incubated with Anti-mouse TRITC antibody. After 2 washes with 0.1% Tween-20, slides were mounted with coverslips using 70% glycerol in 1X PBS. Fibres were imaged using a Nikon A1R HD confocal microscope (Eclipse *Ti-E*, Nikon) at 60X and 1.5 zoom. Between 100 and 250 fibres were measured using ImageJ software from 3 independent experiments and p-values were calculated using Graph Pad Prism software.

### Chromatin fractionation

For chromatin fractionation cells were seeded in 60 mm dishes to a confluency of 30-40%. The cells were synchronised using double thymidine block and transfected as mentioned earlier and collected at 0 hours and 4 hours post G1/S release. The cells were washed with 1X PBS and resuspended in 200 µl of buffer A (10 mM HEPES (pH 7.9), 10 mM KCl, 1.5 mM MgCl_2_, 0.34 M sucrose, 10% glycerol, 1 mM dithiothreitol, and protease inhibitor mixture (Roche Molecular Biochemicals, Switzerland). Triton X-100 was added to a final concentration of 0.1%, and the cells were incubated for 5 min on ice. Nuclei were collected in the pellet (P1) by centrifugation at 1500 X *g*, 4 min, 4°C. The supernatant (S1) was further clarified by centrifugation at 13,000 X *g*, 10 min, 4°C to remove cell debris and insoluble aggregates. This supernatant was designated S2. The nuclei were washed once with buffer A and then lysed in 200 µl of buffer B (3 mM EDTA, 0.2 mM EGTA, 1 mM dithiothreitol, and protease inhibitor mixture). After a 10-min incubation on ice, soluble nuclear proteins (S3) were separated from chromatin by centrifugation (2000 X g, 4 min at 4°C). Isolated chromatin (P3) was washed once with buffer B and spun down at high speed (13,000 X g, 1 min 4°C).

### CldU-chromatin immunoprecipitation (CldU-ChIP)

HeLa cells were synchronised and transfected with siRNA for *BRG1* and/or plasmids for wild type *PLK1* for 48 hours. At the beginning of the second thymidine block, the cells were treated with 100 µM CldU which remained in the media until all the samples were harvested. At the end of the treatment cells were fixed with 1% formaldehyde for 10 min and washed twice with ice-cold 1X PBS. Cell nuclei were isolated using 1 ml of nuclear preparation buffer followed by centrifuged at 12,000 rpm for 2 min at 4 °C. The nuclear pellets were washed with 1 ml of nuclear preparation buffer and lysed using freshly prepared ice-cold sonication buffer-1 (EDTA (pH 8.0): 0.01 M; Tris-HCl (pH 8): 0.05 M; SDS: 1%) for 15 min at 4°C. After incubation, sonication buffer-2 (NaCl: 0.30 M; EDTA (pH 8.0): 0.04 M; Tris-HCl (pH 8): 0.10 M; NP-40 (v/v): 2%; NaF: 0.04 M) was added, and the samples were mixed by vortexing briefly. The nuclear lysate was sonicated in a water bath ultra-sonicator until an average fragment length of 250 bp to 500 bp, and shearing is checked by running samples on 1.5% agarose gel electrophoresis. The supernatant was collected by centrifuging the samples at 12,000 rpm for 10 min at 8°C. The cleared samples were heated at 95 °C to denature the DNA and diluted further by adding 600 µl of dilution buffer. Pre-clearing of diluted samples was done using 50% slurry of Protein G Agarose beads (pre-equilibrated in dilution buffer) at 4°C on a rotating wheel for 1 hour followed by centrifuging at 12,000 g for 2 min to recover the supernatant. The DNA is extracted from chromatin using phenol-chloroform method and quantified using NanoDrop™ 2000 spectrophotometer (Thermo Fisher Scientific, USA).

BRG1 antibody was added in 50 µg chromatin DNA and volume was made up to 200 µl using IP buffer. Antibodies chromatin mix is incubated for 16 hours at 4°C on a rotating wheel. For experimental control, beads alone samples (chromatin without antibody) were used. Protein AG beads were also blocked with BSA (0.1 µg/µl) and salmon sperm DNA (75 ng/µl) in IP buffer (NaCl: 0.15 M; EDTA (pH 8.0): 0.02 M; Tris-HCl (pH 8): 0.5 M; NP-40 (v/v): 1%; NaF: 0.02 M; Sodium deoxycholate (w/v): 0.5%; SDS: 0.1%) for 16 hours at 4°C. After incubation, 30 µl BSA and salmon sperm DNA blocked bead were added to each IP sample and incubated further for 2 hours at 4°C on a rotating wheel. Following incubation, the beads-immunoglobulin-DNA complexes were pelleted by centrifuging at 2,000 rpm for 2 min at 4°C. These complexes were washed with 1 ml of each of the following buffers: ice-cold IP buffer twice, chip wash buffer thrice (LiCl: 0.5 M; EDTA (pH 8.0): 0.02 M; Tris-HCl (pH 8): 0.1 M; NP-40 (v/v): 1%; NaF: 0.02 M; Sodium deoxycholate (w/ v):1%), ice-cold IP Buffer twice, and TE Buffer twice. The samples were mixed with the buffer by inverting the tubes at least 6 times and centrifuging at 2,000 rpm for 2 min at 4°C.

After the last wash, the supernatant was removed as much as possible without disturbing the beads. The DNA was isolated using chelex resins at room temperature. The washed beads were mixed with 100µl of 10% (w/v) chelex 100 slurry (Bio-Rad, USA) prepared in water. The beads were mixed properly with cut-off tips, and the suspension was frequently inverted. The samples were then vortexed briefly for 10 sec and heated at 95°C for 10 min. After cooling down to room temperature, the samples were centrifuged for 1 min at 12000 rpm. Then, 10 µl proteinase K and 10 µl 10 mg/ml RNase were added to each sample. The samples were then incubated at 55°C for 3 hours. Subsequently, proteinase K was inactivated at 95°C for 10 min. The samples were then centrifuged at 12000 rpm for 1 min. About 70-80 µl of supernatant was transferred to a fresh microcentrifuge tube and qRT-PCR was performed.

### ChIP-sequencing library preparation

ChIP-seq library was prepared with 100 ng DNA using NEBNext^®^ Ultra^™^ DNA Library Prep Kit for Illumina^®^ (Catalog # E7370) as per the manufacturer’s instructions. The prepared ChIP DNA and Input DNA libraries were sequenced by Genotypic Technologies, Bengaluru, India.

### ChIP-sequencing data analysis

The data was provided in SangerFASTQ.gz compressed format. The raw reads of ChIP sequencing data (GSE137250) were processed on the Galaxy (https://usegalaxy.org) platform. The raw reads were first trimmed using trimmomatic (version 0.36.5) to remove the adaptor sequence and to improve per base sequence quality. This was followed by quality control analysis using FastQC. The processed reads were aligned to the reference genome (hg38) with the BOWTIE2 tool. The duplicate reads from the aligned reads were identified and removed using Picard Markduplicates in the SAM flags field for each read (<u>http://</u> <u>picard.sourceforge.net/</u>). Duplicate reads were filtered out using the Filter SAM tool (version 1.8). The aligned BAM files for biological duplicates of both BRG1 ChIP sample as well as Input control sample were merged in a single BAM file before peak calling. Peak calling of cleaned reads was done by MACS2 tool (MACS2 Version 2.1.1.20160309.0). ChIP and Input samples were used for background normalization. Binding of BRG1 on genome was visualised using Integrative Genomics Viewer (IGV) tool.

### De novo motif analysis

Using the FASTA format of the identified BRG1 peaks, MEME Suite was performed for motif-based sequence analysis, and the motif with the lowest E-value was selected for further study (Bailey et al., 2009).

### SNS and MCM7 sequencing analysis

SNS-seq data and MCM7 ChIP-seq data were obtained from GSE37757 and GSM2863239, respectively. SNS-seq data files were processed like BRG1 ChIP-seq data and broad peak calling was done using MACS broadpeakcaller to obtain SNS-peaks. MCM-seq data was processed and aligned with hg38 using bowtie 0.12.8 and peak calling was done using macs2 callpeak -t MCM7.bam -c input.bam --name MCM7 --gsize hs --nomodel --extsize 160 --broad --broad-cutoff 5e-2 --to-large --pvalue 1e-3 commands.

### DNaseI-seq, Repli-seq and histone modification data analysis

DNaseI-seq, Repli-seq, H3 histone modification and H4K20me sequencing data were obtained from GSE32970, SRP001393.1, GSE29611 and GSE134988 respectively. The data processing and alignment was done on the Galaxy (https://usegalaxy.org) platform or on the terminal using either BWA or bowtie alignment tools. Post alignment peak calling was done using MACS2 (Version 2.1.1.20160309.0) and bed file processing, in the case of repli-seq, and peak intersection was done using bedtools. The BigWig files of identified peak were visualized with the Integrative Genomics Viewer (IGV) to show the binding of protein at various genomic regions.

### Protein-protein docking

The PBD domain of PLK1 was retrieved from RCSB database (https://www.rcsb.org) with PDB accession 5NN1. Similarly, the BRG1 bromodomain was retrieved using PDB accession number of 2GRC. To dock BRG1 bromodomain with PLK1 PBD, their respective PDB ids were used on ZDOCK under default settings. The resulting docked files were visualised through PyMol.

### Statistical analysis

At least three independent experiments in triplicates were performed for all the qRT-PCR experiments. Statistical significance (*p <0.05; unpaired student’s t-test) was performed using Sigma plot. Unless otherwise stated, all experiments were presented as average ± s.e.m. of three independent biological replicates. *GAPDH* was used as internal control for qRT-PCR experiments and β-actin was used as internal control for western blotting.

## Results

### BRG1 is enriched on dormant replication origins

To investigate the role of BRG1 in DNA replication, ChIP-seq was performed to determine the genome-wide occupancy of BRG1 in HeLa cells. Analysis of the ChIP-seq data revealed that BRG1 was predominantly enriched in intergenic regions characterized by repetitive motifs, notably those containing TCTG and AG dinucleotide repeats (Fig. S1A, B). To determine whether BRG1 localizes to known replication origins, we integrated three datasets: short nascent strand sequencing (SNS-seq), replication timing profiles from Repli-seq, and MCM7 ChIP-seq. SNS-seq identifies replication origins through the enrichment of newly synthesized short DNA fragments, while Repli-seq provides replication timing profiles across S-phase, exhibiting characteristic ‘inverted V’ patterns at replication origins (Fig. S1C). MCM7 ChIP-seq was used to define origins that are actively firing during replication initiation.

Replication origins were classified into firing and dormant categories by overlapping SNS-seq peaks with MCM7-bound sites, using established criteria. SNS-seq analysis identified 150,528 putative origin peaks, while MCM7 ChIP-seq identified 77,808 binding sites. Intersecting these datasets yielded 21,220 firing origins (Fig. 1A). The remaining 129,308 SNS-seq peaks that did not overlap with MCM7 sites were designated as dormant replication origins. We next assessed BRG1 localization at replication origins by intersecting BRG1 ChIP-seq peaks with the SNS-seq dataset. Of the 15,830 BRG1-bound regions, 13,970 (88.3%) overlapped with SNS-seq peaks, indicating BRG1 is associated with replication origin sites (Fig. 1B). To further differentiate whether these BRG1-occupied origins were firing or dormant, we compared the 13,970 BRG1-bound origin peaks with the 21,220 firing origins. Only 111 (0.007%) of BRG1-bound sites overlapped with firing origins. In contrast, 6,375 of the 13,970 BRG1-bound sites (45.63%) overlapped with dormant origins (Fig. 1C, D). Together, these data demonstrate that BRG1 is predominantly enriched at dormant replication origins, suggesting a potential role in regulating replication origin licensing or activation under stress conditions.

**Figure 1:**
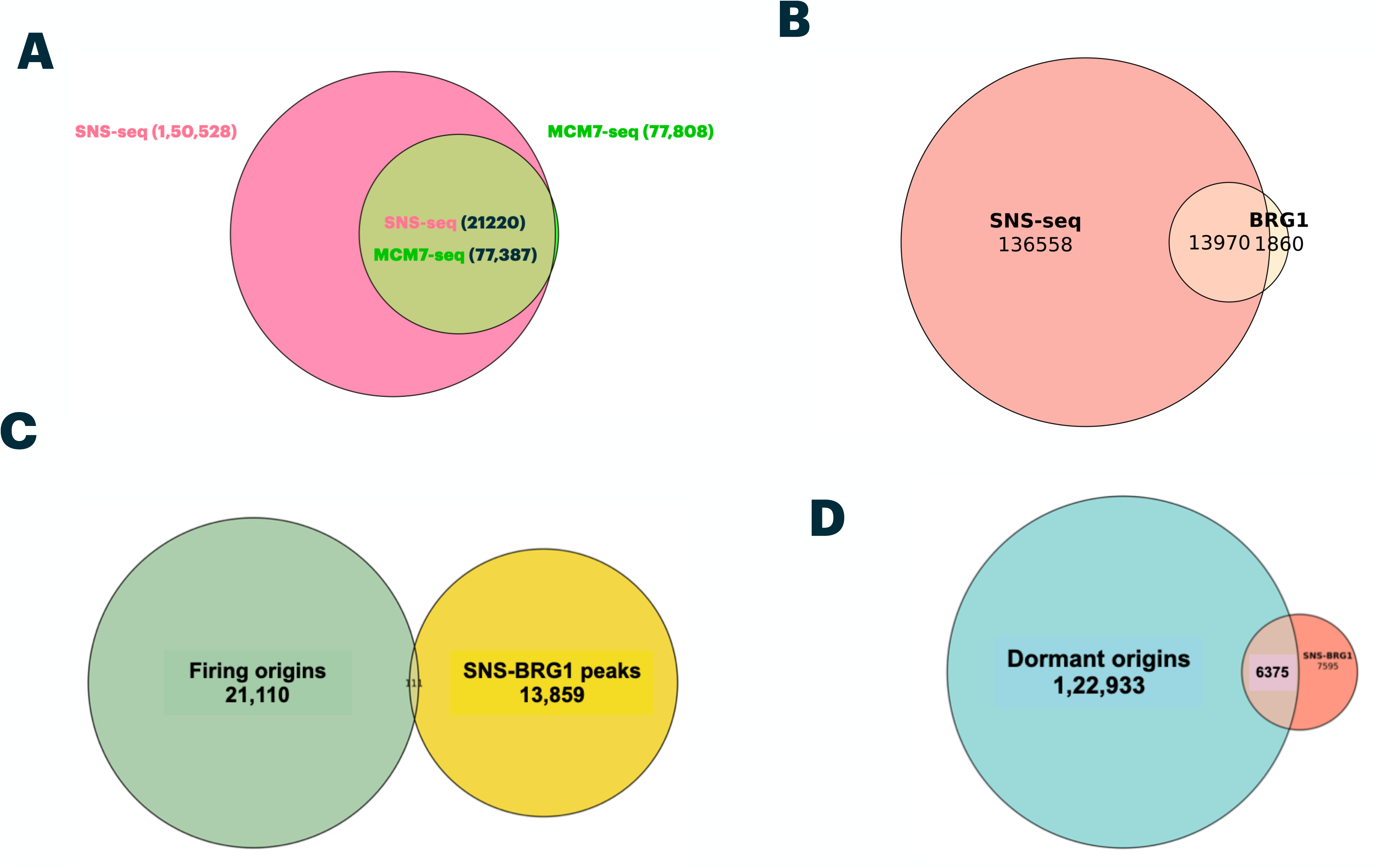
BRG1 is enriched on dormant replication origins. Overlap between (A) SNS-seq and MCM7 ChIP-seq data, (B) SNS-seq and BRG1 ChIP-seq, (C) Firing origins, obtained from SNS-MCM7 intersected regions, with SNS-BRG1 (D) Dormant origins, obtained from regions outside of SNS-MCM7, intersection with BRG1-SNS regions.

#### BRG1 occupied dormant replication origins are present on late replicating heterochromatin regions

To further characterize the replication timing of BRG1-bound origins, the S50 values from Repli-seq data was estimated. The S50 metric represents the time point during S-phase at which 50% of a given genomic region has been replicated, with values ranging from 0 (early) to 1 (late). BRG1-bound firing origins exhibited a median S50 value of 0.45, indicating early to mid-S phase replication, while BRG1-bound dormant origins displayed a higher median S50 value of 0.60, consistent with mid-to-late S-phase replication (Fig. 2A). Given previous observations that firing origins tend to be GC-rich whereas dormant origins are GC-poor, the GC content of BRG1-bound firing (n=111) and dormant (n=6,375) origins were compared. Consistent with expectations, the firing origins exhibited significantly higher GC content compared to dormant origins (Fig. 2B). To assess the sequence features commonly associated with replication origins, including G-quadruplex (G4) motifs, CpG islands, and chromatin accessibility, we analyzed the potential for G4 formation using the QGRS Mapper tool. BRG1-bound firing origins were enriched for G4-forming sequences (91 G4 motifs), whereas dormant origins contained substantially fewer G4 motifs (50) (Fig. 2C).

**Figure 2:**
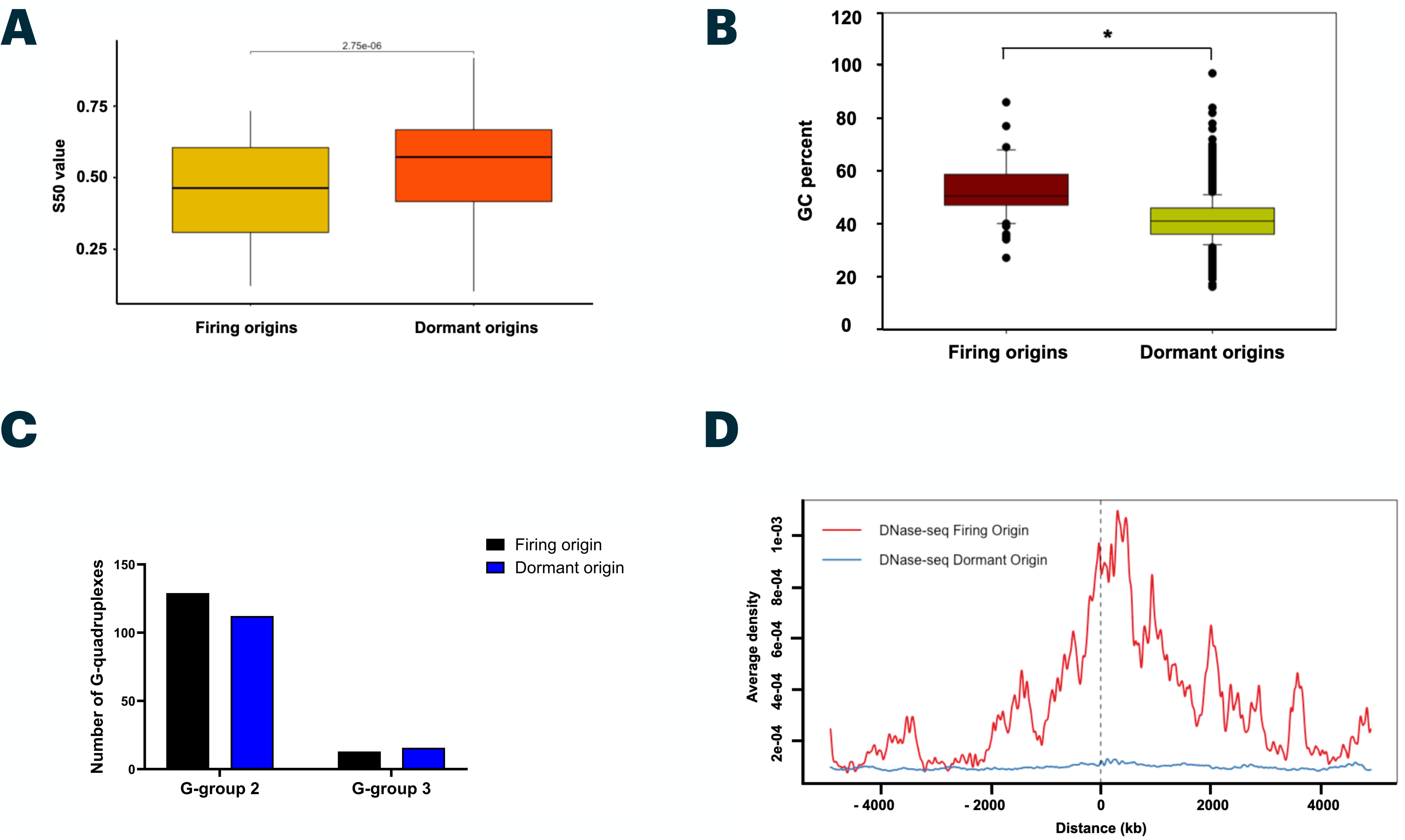
BRG1 occupied dormant replication origins are present on late replicating heterochromatin regions. Distribution of S50 values for firing and dormant origins ranging from 0 (early) to 1 (late) replication time (A). Quantification of (B) GC percent from 111 firing and 6375 dormant origin peaks (C) G-quadruplex forming propensity as measured by QGRS tool (D) Aggregation plots of DNase-seq peaks surrounding firing and dormant origins. The statistical significance (*p <0.05) was calculated using unpaired student’s t-test. The data in panel A is presented as mean GC percentages through Sigma plot.

Next, chromatin accessibility and epigenetic marks around BRG1-bound origins was evaluated. DNase I hypersensitivity analysis revealed that firing origins were more accessible, with a pronounced hypersensitivity signal surrounding BRG1-bound firing origins relative to dormant sites (Fig. 2D). Enrichment of active chromatin marks, including H3K4me2, H3K4me3, H3K9ac, and H3K27ac, was also greater at firing origins than at dormant ones (Fig. S2A, B), consistent with their localization near transcriptionally active genomic regions. We then analyzed heterochromatic histone modifications commonly associated with late replication. BRG1-bound dormant origins, but not firing origins, showed co-enrichment with repressive histone marks H3K9me3 and H3K27me3 (Fig. S2C). Furthermore, assessment of H4K20 methylation states showed that BRG1-occupied regions overlapped with H4K20me3, a mark associated with late-replicating heterochromatin and dormant origins, but not with H4K20me1, which is implicated in pre-replication complex (pre-RC) formation (Fig. S2D). Collectively, these data indicate that BRG1-bound dormant replication origins are enriched in late-replicating, heterochromatic regions characterized by low GC content, reduced chromatin accessibility, and repressive histone modifications.

#### BRG1 regulates replication dynamics at dormant replication origins

To experimentally validate the *in silico* findings, the localization of BRG1 at active replication sites was assessed in HeLa cells using immunofluorescence-based detection of IdU incorporation. Cells transfected with either control siRNA (siC) or *BRG1*-targeting siRNA (si*BRG1*) were synchronized using a double thymidine block and collected 4 hours post-release, corresponding to mid-to-late S phase. Co-localization analysis revealed that BRG1 significantly overlapped with IdU signals in siC-treated cells (Pearson’s correlation coefficient = 0.32), while this overlap was significantly reduced in BRG1-depleted cells (Pearson’s coefficient = 0.21, *p* < 0.001), suggesting diminished BRG1 presence at replication sites (Fig. 3A–C). To confirm BRG1 occupancy at replication origins, ChIP-qPCR was performed at selected genomic loci classified as either firing (*TOP1, CTNNA3, STX6, SNX16*) or dormant (*RBM39, MET, CDK14, CAV2, PREX2, CUL2, IL1R2, CD40*) origins, based on SNS-seq and MCM7 ChIP-seq analysis. Consistent with ChIP-seq data, BRG1 was enriched at nearly all selected origins, except *TOP1* and *RBM39*, at the G1/S boundary (0 h). At 4 hours post-release, BRG1 occupancy increased at all examined loci except *TOP1*, *MET*, and *PREX2* (Fig. S3A). This increase in BRG1 binding correlated with elevated CldU incorporation at most origins—excluding *TOP1* and *LMNB2*—indicating enhanced replication activity (Fig. S3B). These data suggest that BRG1 occupancy at replication origins increases as S-phase progresses. To evaluate the functional impact of BRG1 on replication dynamics, DNA fiber assays were performed in siC- and si*BRG1*-transfected HeLa cells synchronized by double thymidine block and released into S-phase. At 4 hours post-release, DNA synthesis as measured by IdU/CldU incorporation was significantly impaired in si*BRG1* cells compared to siC controls, consistent with previous reports (Fig. 4A). Notably, the inter-origin distance was reduced in *BRG1*-depleted cells (Fig. 4B, C), indicating activation of nearby dormant origins in response to compromised replication. We next examined whether BRG1 is differentially required for replication through firing versus dormant origins. CldU incorporation was elevated on firing origins in both siC and si*BRG1* cells, indicating that BRG1 is dispensable for replication through these sites. In contrast, CldU incorporation was significantly reduced on dormant origins in siBRG1 cells, suggesting that BRG1 is essential for replication from these loci (Fig. 4D). Together, these results indicate that while BRG1 is present at both firing and dormant origins, it is specifically required for efficient replication from dormant replication origins.

**Figure 3:**
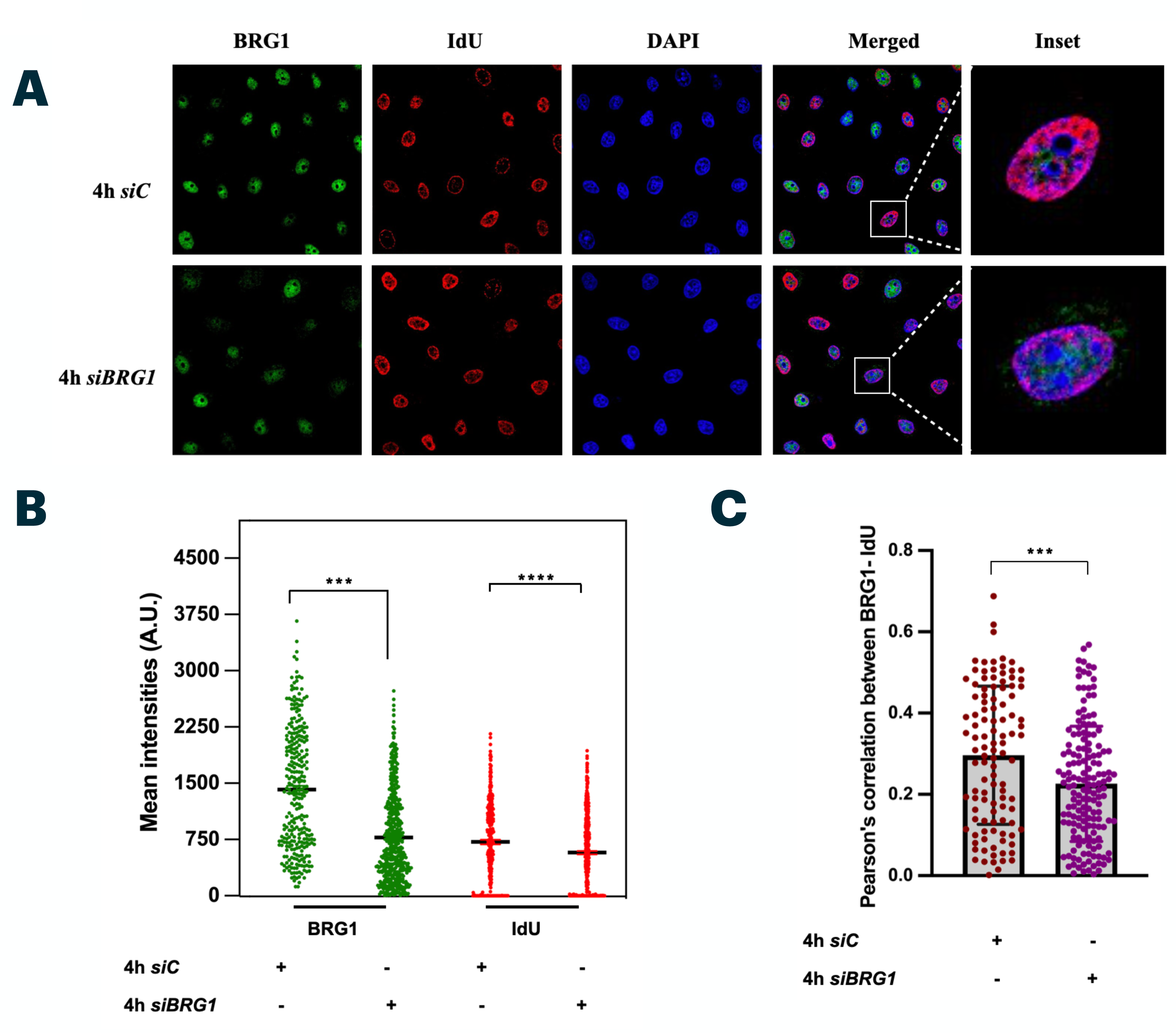
BRG1 regulates replication dynamics at dormant replication origins. (A) Co-localization of BRG1 and IdU was monitored in control and *BRG1* depleted S phase synchronised cells. (B) Quantification of FITC (BRG1) and TRITC (IdU) nuclear intensities. (C) Measurement of persons co-efficient of corrleation between BRG1 and IdU in siC and si*BRG1* cells. The statistical significance (*p <0.05) was calculated using an unpaired student’s t-test. For confocal image quantification, n > 200 cell nuclei were taken from three independent biological replicates.

**Figure 4:**
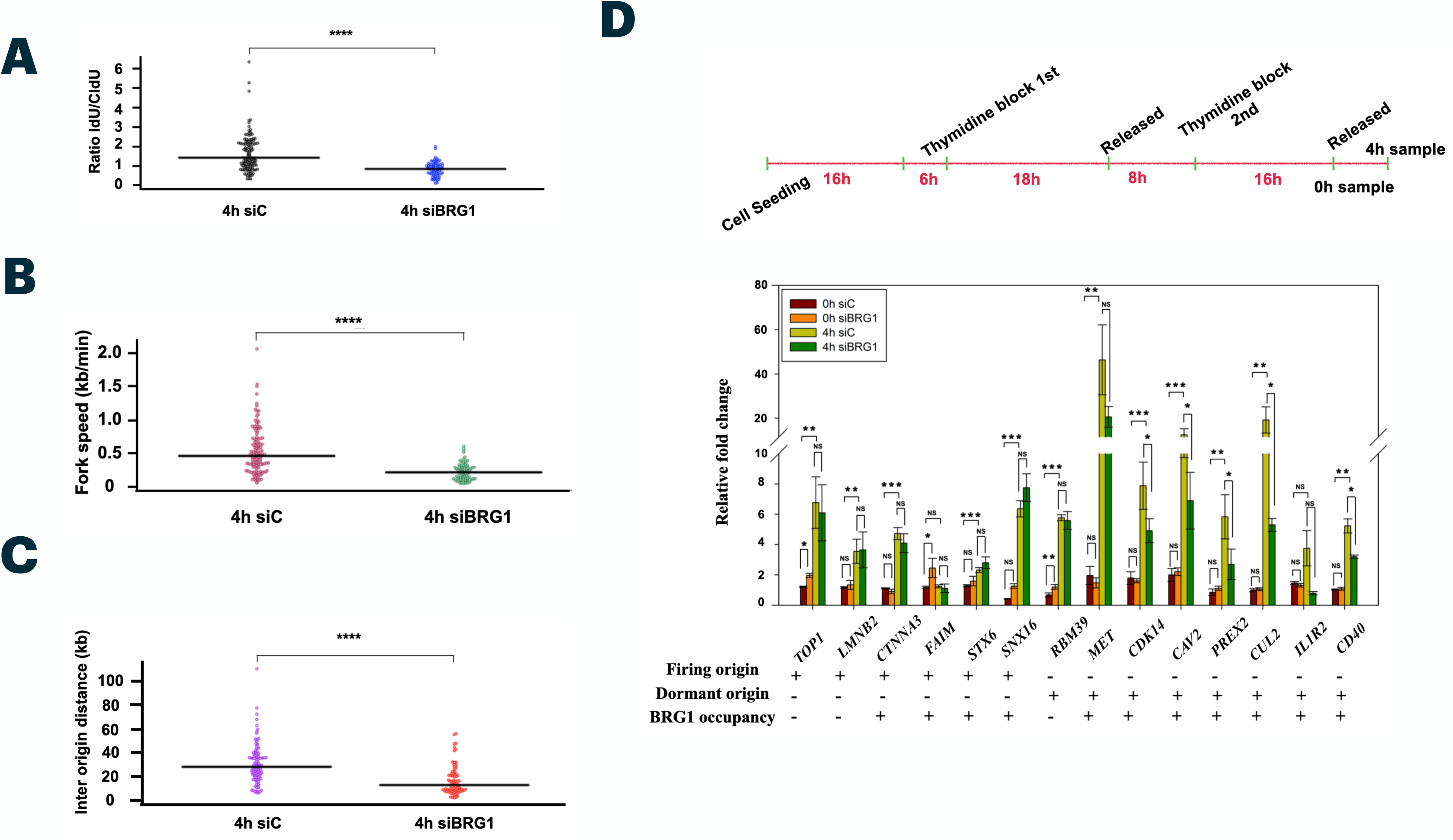
Replication dynamics in *BRG1* depleted HeLa cells. (A-D) DNA fibre analysis of S phase synchronised HeLa cells transfected with siC or *siBRG1* followed by sequential labelling with CldU and IdU. (A) The mean distribution of the IdU/CldU incorporation ratio was calculated by measuring the length (in µm) of the IdU labelled track (red) and CldU labelled track (green). (B) Quantification results of fork speed and (C), inter-origin distances. (D) Replication progression through selected origins in *BRG1* depleted cells at 0 hours and 4 hours post release from double thymidine block. Replication progression through indicated origins was measured by immunoprecipitation of CldU in siC and si*BRG1* cells labelled with CldU for 16 hours and 18 hours for 0 hours and 4 hours respectively.. The swarm plot shows the mean distribution and statistical significance (*p <0.05) was calculated using an unpaired student’s t-test. For quantification n> 200 DNA fibres were taken from three independent biological replicates.

#### BRG1 is required for PLK1 foci formation during S-phase

Given the role of PLK1 in phosphorylating Orc2 and regulating dormant origin activation under replication stress, we next investigated whether BRG1 functionally interacts with PLK1 to modulate replication dynamics. Using an in silico docking approach, we evaluated the potential interaction between the BRG1 bromodomain (PDB ID: 2GRC) and the PLK1 polo-box domain (PBD, PDB ID: 1Q4O). Docking analysis using ZDOCK revealed a moderate interaction involving four polar contacts: PLK1 residues Arg-158, Arg-313 (repeatedly), and Arg-324 interact with BRG1 residues Glu-1499, Glu-1493, Glu-1496, and Glu-1542 (Fig. S4A). This predicted interaction was experimentally supported by co-immunoprecipitation in synchronized HeLa cells, demonstrating a physical association between BRG1 and PLK1 (Fig. 5A). To determine whether BRG1 regulates PLK1 expression, we examined PLK1 transcript and protein levels in siBRG1 versus siC cells in asynchronous and at 0 and 4 hours post-release from double thymidine block. Quantitative PCR showed no significant change in PLK1 mRNA levels upon BRG1 depletion (Fig. 5B). However, PLK1 protein expression was markedly reduced in siBRG1 cells (Fig. S4B and 5C), indicating post-transcriptional regulation, potentially through reduced stability or translation efficiency. Confocal microscopy analysis of BRG1 and PLK1 localization revealed a notable reduction in the number and intensity of PLK1 foci in siBRG1 cells during S-phase (Fig. S4C, Fig. 5D), suggesting BRG1 may be necessary for PLK1 chromatin association. To confirm this, cytoplasmic and chromatin-bound protein fractions were isolated from synchronized HeLa cells. Western blot analysis demonstrated reduced levels of both BRG1 and PLK1 in the chromatin-bound fraction of siBRG1 cells compared to controls (Fig. 5E). Together, these findings suggest that BRG1 directly or indirectly interacts with PLK1 and is required for proper localization and accumulation of PLK1 on chromatin during S-phase, which may in turn influence replication dynamics, particularly at dormant origins.

**Figure 5:**
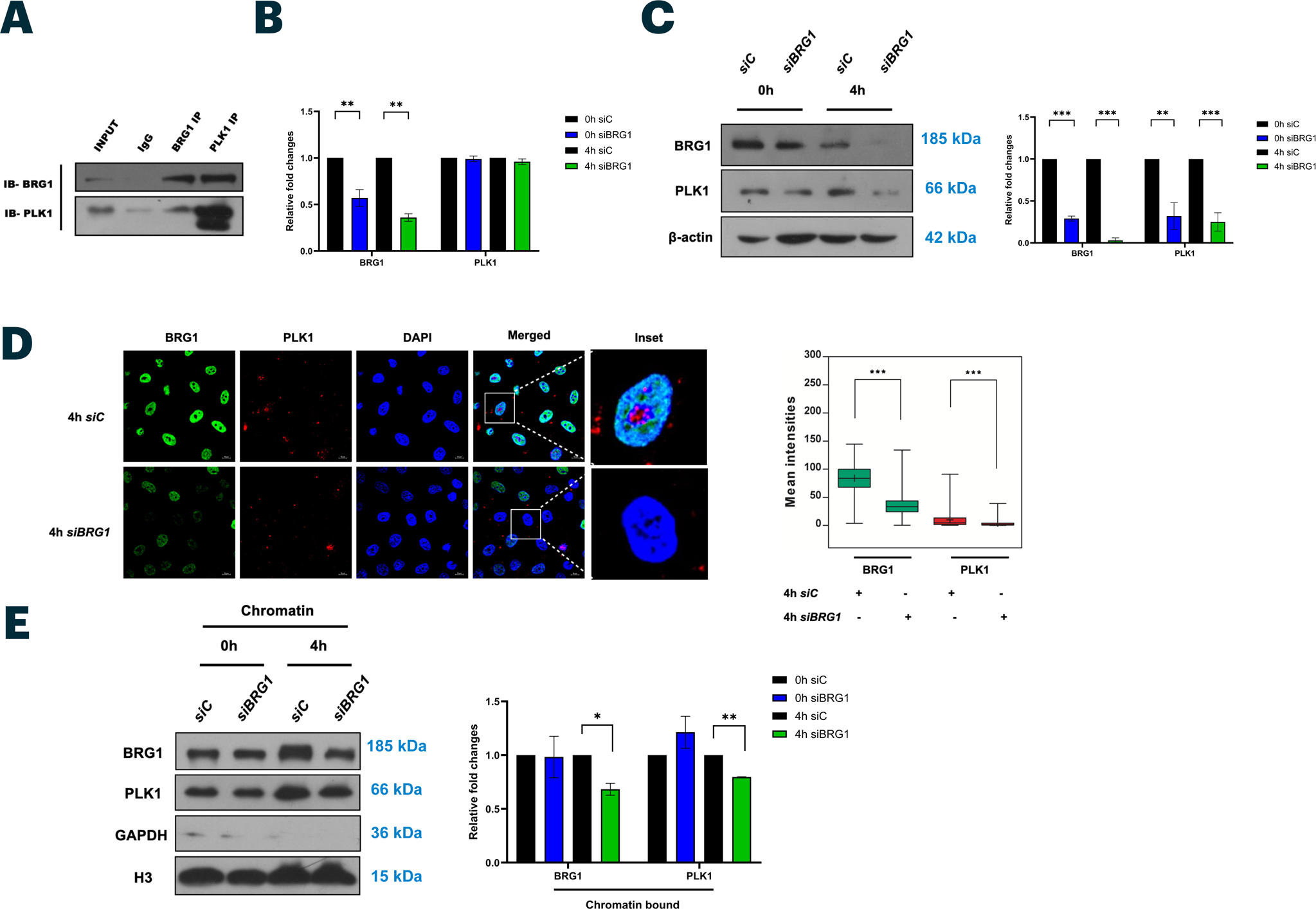
BRG1 is required for PLK1 foci formation during S-phase. (A) Co-immunoprecipitation of BRG1 and PLK1 in HeLa cells. mRNA (B) and protein (C) expression of BRG1 and PLK1 in control (siC) and BRG1 depleted (*siBRG1) G1*/S and S phase synchronised cells.(D) Mean nuclear intensities of BRG1 and PLK1 protein in S-phase synchronised siC and *siBRG1* cells. (E) Cytosolic and chromatin bound protein levels of BRG1, PLK1, GAPDH (cytosolic control) and H3 (nuclear control) in siC and si*BRG1* mid-S phase synchronised cells. The statistical significance (*p <0.05) was calculated using an unpaired student’s t-test. All experiments presented as average ± SEM of three independent biological replicates. GAPDH was used as an internal control for qRT-PCR experiments and β-actin was used as an internal control for western blotting. The intensities of western blots were quantitated using Image J software.

#### PLK1 stabilises BRG1 protein

To investigate the regulatory relationship between BRG1 and PLK1, wild-type PLK1 was overexpressed in synchronised HeLa cells. PLK1 overexpression led to increased BRG1 protein levels at both 0h and 4 hours post thymidine block release, indicating that PLK1 positively regulates BRG1 protein expression (Fig. 6A). To determine whether this regulation occurs at the transcriptional or post-transcriptional level, both BRG1 and PLK1 transcript and protein levels were quantified in si*BRG1*-treated cells overexpressing wild-type PLK1 under the same synchronization conditions. Although *BRG1* mRNA remained reduced in si*BRG1* cells, PLK1 overexpression significantly elevated BRG1 protein levels (Fig. 6B). We further performed immunofluorescence in si*BRG1* cells over expressing wild type PLK1 and quantified nuclear intensities of both BRG1 and PLK1. Consistent with the protein levels, nuclear levels of BRG1 were significantly increased in si*BRG1* cells overexpressed with wild type PLK1 as compared to control (Fig. S5A). The concurrent increase in both PLK1 mRNA and protein confirmed effective PLK1 overexpression (Fig. S5B, 6B). These findings suggest that PLK1 regulates BRG1 expression post-transcriptionally, possibly by enhancing BRG1 protein stability or translation efficiency. To directly assess whether PLK1 activation is necessary for BRG1 protein stability, a PLK1 phosophorylation inactive mutant, T210A, where threonine 210 was substituted with alanine, was generated and validated by sequencing. This mutant was overexpressed in si*BRG1* cells, and BRG1 and PLK1 protein levels were subsequently assessed. While the T210A mutant showed elevated PLK1 protein levels at both synchronization time points, BRG1 levels were notably reduced compared to controls (Fig. 6C). These results indicate that PLK1 kinase activity, rather than its mere presence, is required for maintaining BRG1 protein stability. To further validate this conclusion, a PLK1 deletion construct lacking the catalytic domain (encompassing both the kinase domain and the T210 activation site) was overexpressed in parental HeLa cells. The PLK1 detection antibody used in this experiment recognizes an epitope within the Polo-box domain (PBD), allowing detection of both full-length and truncated PLK1 proteins. Interestingly, overexpression of the PLK1 deletion construct also elevated endogenous PLK1 levels at both synchronization points (Fig. 6D). However, consistent with previous findings, BRG1 protein levels were reduced in these cells, supporting the conclusion that PLK1’s catalytic activity is essential for BRG1 protein stability.

**Figure 6:**
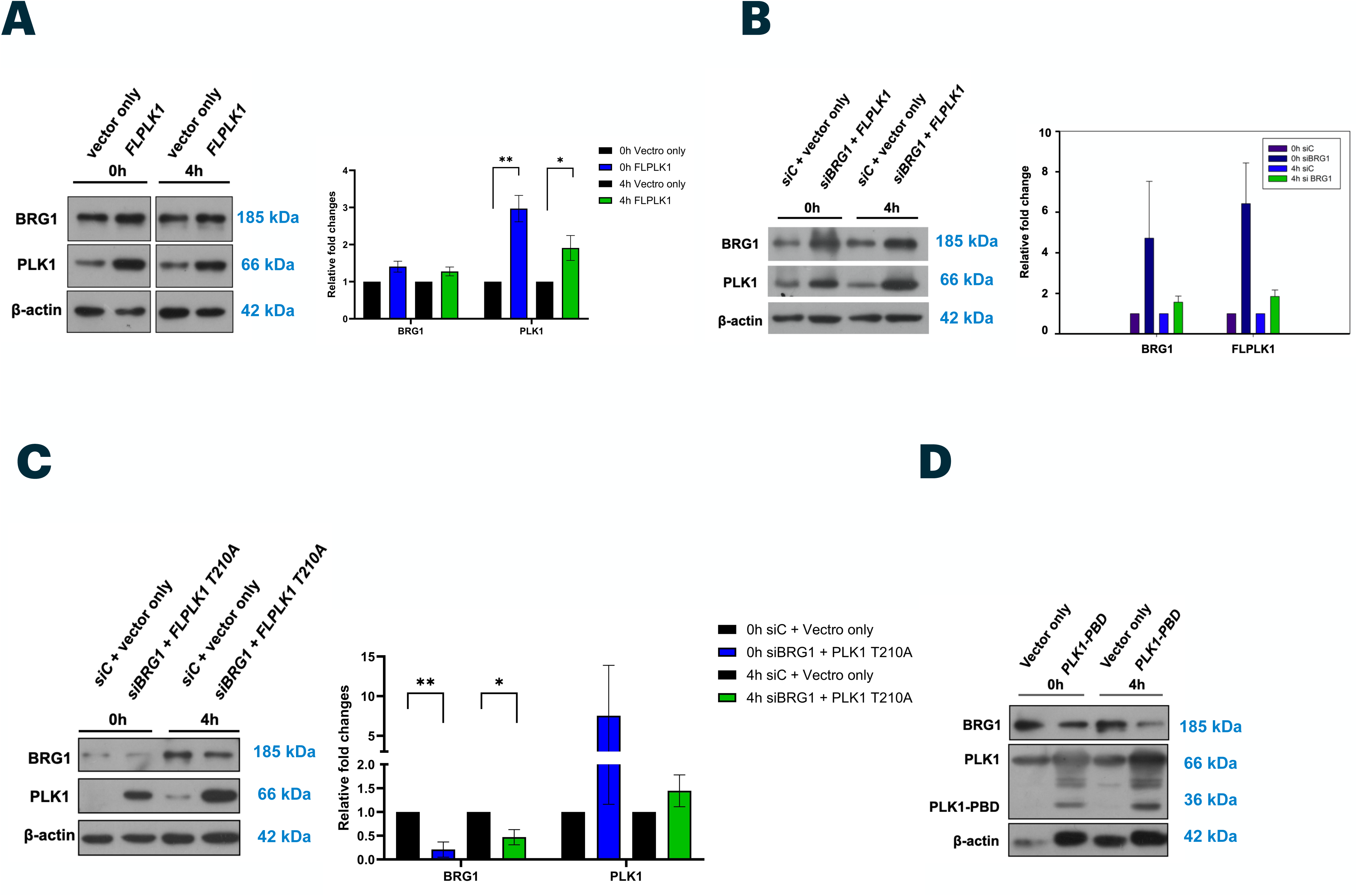
PLK1 stabilises BRG1 protein. Protein expression of BRG1 and PLK1 in control and wild type PLK1 overexpressed (A) control and BRG1 depleted (siBRG1) wild type PLK1 overexpressed (B) and control and BRG1 depleted mutant PLK1T210 (C) overexpressed cells at 0h and 4 hours post-release from double thymidine block. (D) Protein levels of BRG1, endogenous wild PLK1 and catalytic domain deleted PLK1 (PLK1-PBD) protein at 0h and 4 hours post release from double thymidine block. The statistical significance (*p <0.05) was calculated using an unpaired student’s t-test. All experiments presented as average ± SEM of three independent biological replicates except for (C) and (D) where n=2. The intensities of western blots were quantitated using Image J software.

#### PLK1 overexpression rescues replication defects in BRG1-depleted cells

To assess the functional interplay between BRG1 and PLK1 in regulating DNA replication, we tested whether overexpression of wild-type PLK1 could rescue replication-associated defects observed in BRG1-depleted cells. HeLa cells were transfected with si*BRG1* and simultaneously overexpressed full-length PLK1. Immunofluorescence microscopy confirmed that BRG1 levels on chromatin were significantly restored in PLK1-overexpressing, BRG1-depleted cells compared to siBRG1 cells transfected with empty vector or control siRNA (siC) (Fig. S5A). We next asked whether PLK1 overexpression could functionally compensate for BRG1 loss during DNA replication. To this end, synchronized cells depleted of BRG1 and overexpressing PLK1 were pulse-labeled with thymidine analogs CldU and BrdU for 20 and 30 minutes, respectively. DNA fiber analysis revealed that PLK1 overexpression restored replication fork speed, DNA fiber length, and inter-origin distances to near-normal levels, effectively rescuing the replication defects caused by BRG1 depletion (Fig. 7E–G). Finally, to determine whether PLK1 overexpression could restore replication origin activity at specific genomic sites, CldU-ChIP-qPCR was performed. This analysis showed increased CldU incorporation at selected replication origins in BRG1-depleted cells overexpressing PLK1, supporting the conclusion that PLK1 can functionally compensate for BRG1 in promoting origin firing, particularly at dormant origins (Fig. 7H). Collectively, these results demonstrate that PLK1 overexpression restores BRG1 protein levels and rescues replication progression in BRG1-depleted cells, highlighting a functional interplay between BRG1 and PLK1 in the regulation of DNA replication dynamics.

**Figure 7:**
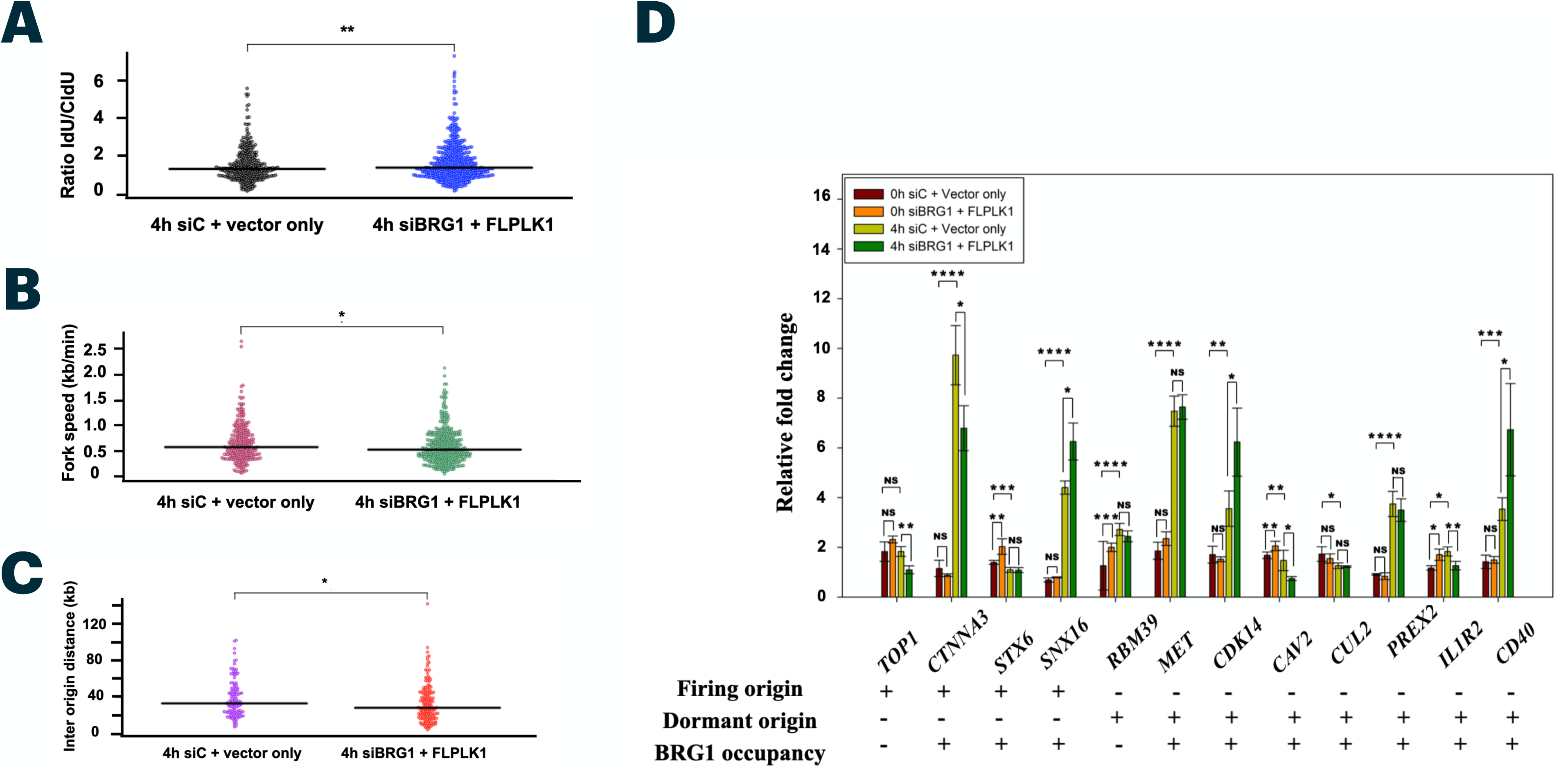
Replication dynamics after overexpression of full length *PLK1* in *BRG1* depleted HeLa cells. DNA fibre analysis of HeLa cells transfected with siC and vector only, and *siBRG1* and full length *PLK1* plasmid. (A) Mean distribution of the IdU/CldU incorporation ratio calculated by measuring the length (in µm) of IdU labelled track (red) and CldU labelled track (green). (B) Quantification results of fork speed and (C) Inter-origin distances. (D) Replication progression through selected origins in control (siC+ vector only) and si*BRG1* PLK1 overexpressed cells at 0 hours and 4 hours post release from double thymidine block. Replication progression through indicated origins was measured by exposing cells to CldU for 16 hours and 18 hours corresponding to 0 hours and 4 hours after double thymidine block respectively followed by chromatin immunoprecipitation.The swarm plot shows the mean distribution and statistical significance (*p <0.05) was calculated using an unpaired student’s t-test. For quantification n > 300 DNA fibres were taken from three independent biological replicates. The statistical significance (*p <0.05) was calculated using unpaired student’s t-test. The data bars are presented as average ± SEM of three independent biological replicates.

## DISCUSSION

A significant portion of the human genome is gene-poor, repetitive, and transcriptionally inactive. Nucleosome assembly and DNA replication must occur across the entire genome—both euchromatic and heterochromatic regions—necessitating chromatin remodeling to ensure accessibility. Replication origins are frequently located in gene-rich, transcriptionally active regions, yet can also be initiated from intergenic and heterochromatic domains. Accessibility to these regions is mediated by ATP-dependent chromatin remodelers such as BRG1 (SMARCA4), which binds chromatin via acetylated histones and is enriched at promoters, enhancers, and other regulatory elements. Previous studies have shown that BRG1 interacts with replication factors and is essential for replication fork progression and origin activation in cells [37, 42–44]. However, its role in the regulation of dormant replication origins remained unexplored.

In this study, we integrated genome-wide BRG1 ChIP-seq data with replication origin maps derived from SNS-seq, Repli-seq, and MCM7 ChIP-seq to demonstrate that BRG1 is preferentially enriched at dormant replication origins, particularly those located in late-replicating, heterochromatic regions. Approximately 43% of BRG1 binding sites were intergenic, and motif analysis revealed enrichment for repetitive DNA elements such as TG and TC repeats—sequences known to impede replication fork progression and induce replication stress [62]. This is consistent with previous findings implicating BRG1 in the formation of Z-DNA structures at repetitive motifs [63], and its affinity for transcription factor motifs such as Smad and Pitx [64, 65].

The intersection analysis showed that 65% of BRG1-bound regions overlapped with dormant origins, while only 7% overlapped with firing origins. Chromatin and sequence features further distinguished these two classes: firing origins were enriched for high GC content, DNase hypersensitivity, G-quadruplex motifs, and active histone marks (e.g., H3K4me2/3, H3K9ac, H3K27ac), consistent with open chromatin and early replication timing. Conversely, BRG1-bound dormant origins exhibited lower GC content, reduced chromatin accessibility, and enrichment for repressive marks such as H3K9me3, H3K27me3, and H4K20me3—hallmarks of compact, late-replicating chromatin. These findings support a model in which BRG1 helps regulate origin activity in repressed chromatin contexts, particularly under conditions that require the activation of dormant origins. Repli-seq analysis confirmed that these BRG1-bound dormant origins replicate primarily during mid-to-late S phase, aligning with their heterochromatic localization. To validate the computational findings, we performed functional experiments in HeLa cells. Immunofluorescence analysis revealed BRG1 localization to replication foci during S phase, which was reduced upon BRG1 knockdown. DNA fiber assays demonstrated impaired fork progression, reduced replication speed, and increased inter-origin distances in BRG1-depleted cells—indicative of replication stress and compensatory origin firing. Notably, replication at dormant origins was significantly compromised, while firing origins were largely unaffected, underscoring the selective role of BRG1 in dormant origin activation.

Previous studies have linked only a few factors such as SIRT1 and FANCI to dormant origin regulation, typically in response to replication stress [66–68]. Our data identify BRG1 as a novel regulator of dormant origins, acting at the interface of chromatin remodeling and replication control. To explored the molecular mechanisms underlying this regulation, we focused on the mitotic kinase PLK1, which is known to regulate pre-replication complex stability and dormant origin activation during replication stress [52, 55–57]. BRG1 depletion reduced PLK1 protein levels without affecting *PLK1* mRNA, suggesting post-transcriptional regulation. Subcellular fractionation revealed diminished chromatin-bound PLK1 in BRG1-depleted cells, implicating BRG1 in PLK1 stabilization or chromatin recruitment during S phase. Strikingly, overexpression of PLK1 in BRG1-depleted cells restored BRG1 protein levels, chromatin occupancy, and replication dynamics, including fork speed and origin activation. These findings suggest a feedback mechanism in which BRG1 and PLK1 co-regulate each other at the protein level to maintain replication fidelity, particularly at dormant origins. In silico docking predicted an interaction between the BRG1 bromodomain and PLK1 polo-box domain, which was confirmed by co-immunoprecipitation, indicating a physical or functional association. Importantly, PLK1 overexpression rescued replication defects caused by BRG1 depletion, even in the absence of PLK1 kinase activity, suggesting that PLK1 may act as a scaffold or stabilizer for BRG1 rather than solely as a kinase in this context. Conversely, overexpression of BRG1 increased PLK1 protein levels, further supporting their reciprocal regulation.

In summary, this study provides the first evidence that BRG1 preferentially regulates dormant origin activation in heterochromatin during DNA replication. BRG1 stabilizes replication through its interaction with PLK1, and loss of either factor leads to impaired replication fork progression, defective origin firing, and replication stress. This mutual regulation suggests a coordinated mechanism to ensure replication completion in transcriptionally repressive regions.

### Future Directions

Several mechanistic questions remain. It is unknown whether BRG1 affects the phosphorylation status of MCM2 and function of RecQL4, a helicase subunit recently linked to dormant origin control [68, 69]. Additionally, BRG1 may influence PLK1 stability via the ubiquitin-proteasome pathway or through acetylation-dependent mechanisms. Whether BRG1’s chromatin remodeling activity (via its ATPase domain) is necessary for dormant origin regulation, and whether PLK1 kinase activity directly modulates BRG1 function or stability, are also important avenues for future investigation. Furthermore, BRG1 has been shown to regulate *ATR* expression—a key component of the replication stress response—and ATR activity is tightly linked to PLK1-mediated checkpoint recovery. Dissecting how BRG1-PLK1 crosstalk integrates with ATR/Chk1 signaling may reveal new insights into replication stress tolerance and genome stability maintenance in heterochromatin.

## AUTHOR CONTRIBUTIONS

Conceptualization, R.M., S.H.; Methodology, R.M., S.H.; Investigation, S.H. D.B.; Resources-R.M.; Writing-original draft, R.M. and S.H.; Writing-review and editing, R.M., S.H. and D.B. Funding acquisition, R.M.; Supervision, R.M.

## ACKNOWLEDGEMENTS

The authors would like to thank the Central Instrumentation facility, School of Life Sciences for confocal microscope and FACS facility. The authors would also like to thank Dr. Sarika Gupta for technical support.

---

**Figure S.1:**
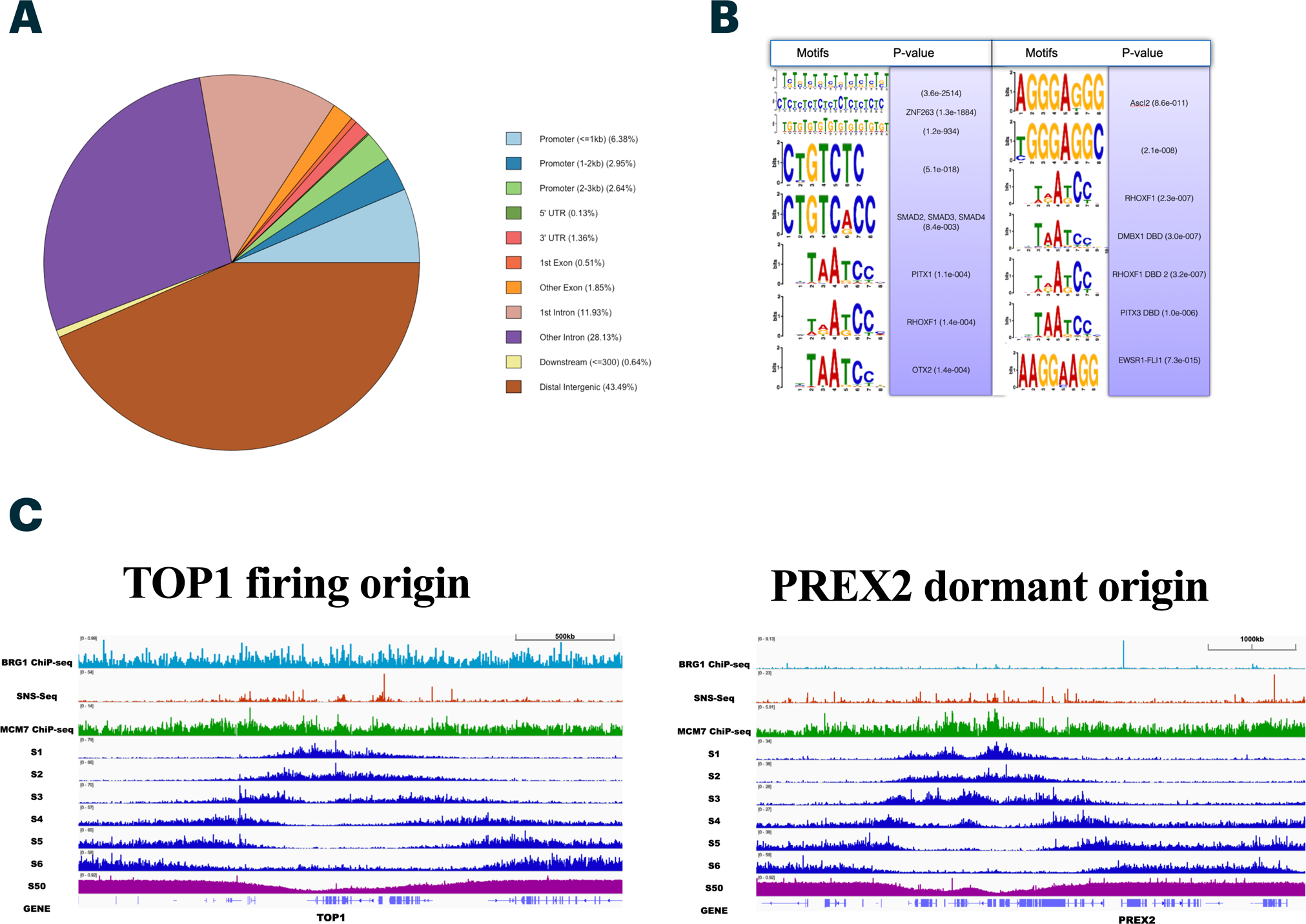
Pie chart of BRG1 genome-wide occupancy. (A) and (B) DNA motifs occupied by BRG1. (C) IGV screen shot showing overlap among BRG1 ChIP-seq, SNS-seq and Repli-seq data on TOP1 (firing) and PREX2 (dormant) origins.

**Figure S.2:**
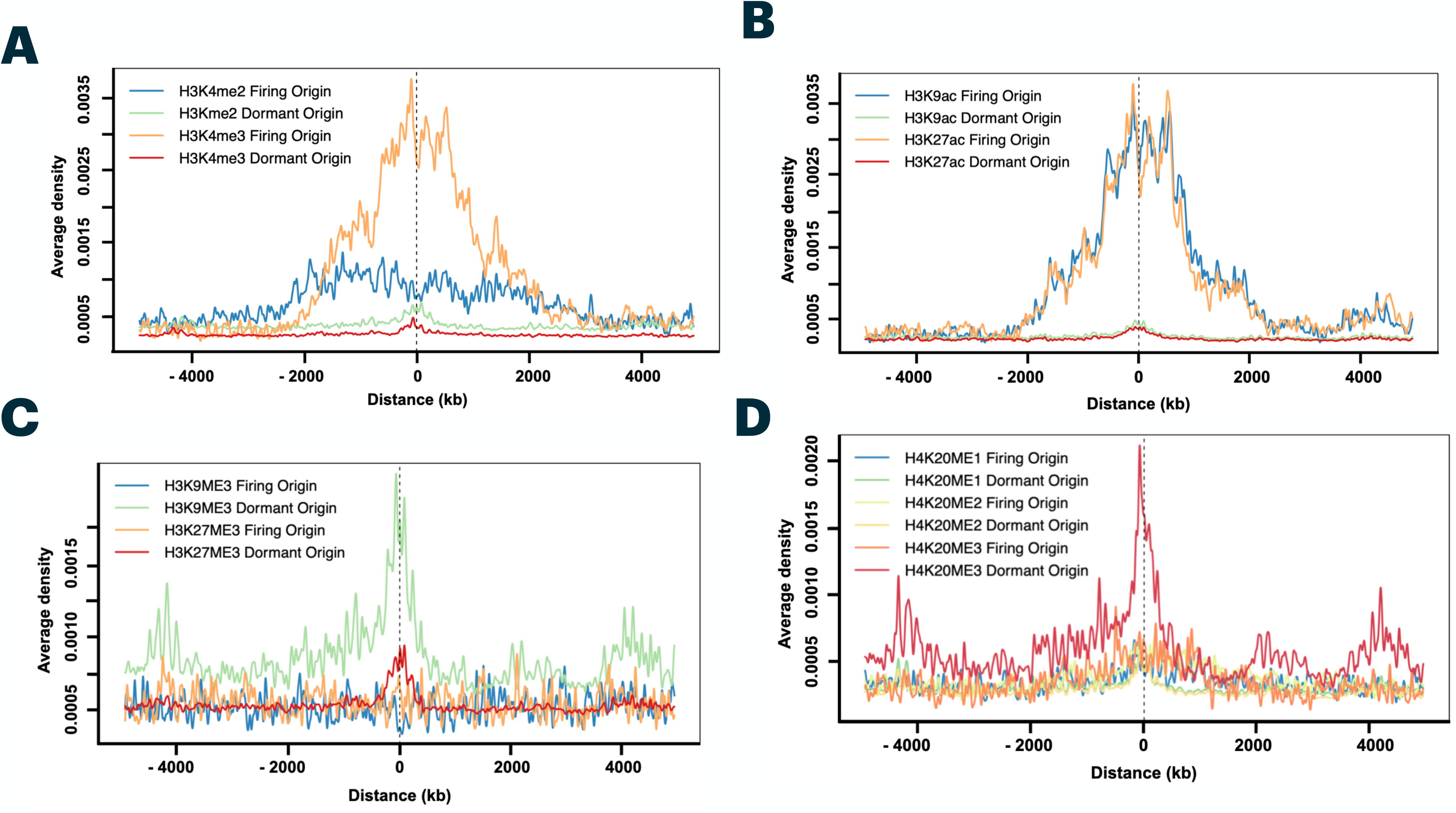
Firing but not dormant origins are enriched in open chromatin histone marks. (A and C) Aggregation plots showing histone modification densities surrounding the indicated origin classes (left). (B and D) Aggregation plots of repressive histone modification densities surrounding the indicated origin classes. Histone modification ChIP-seq signals around ±5 kb from the centre of each BRG1 occupied firing and dormant origins peak are also shown.

**Figure S.3:**
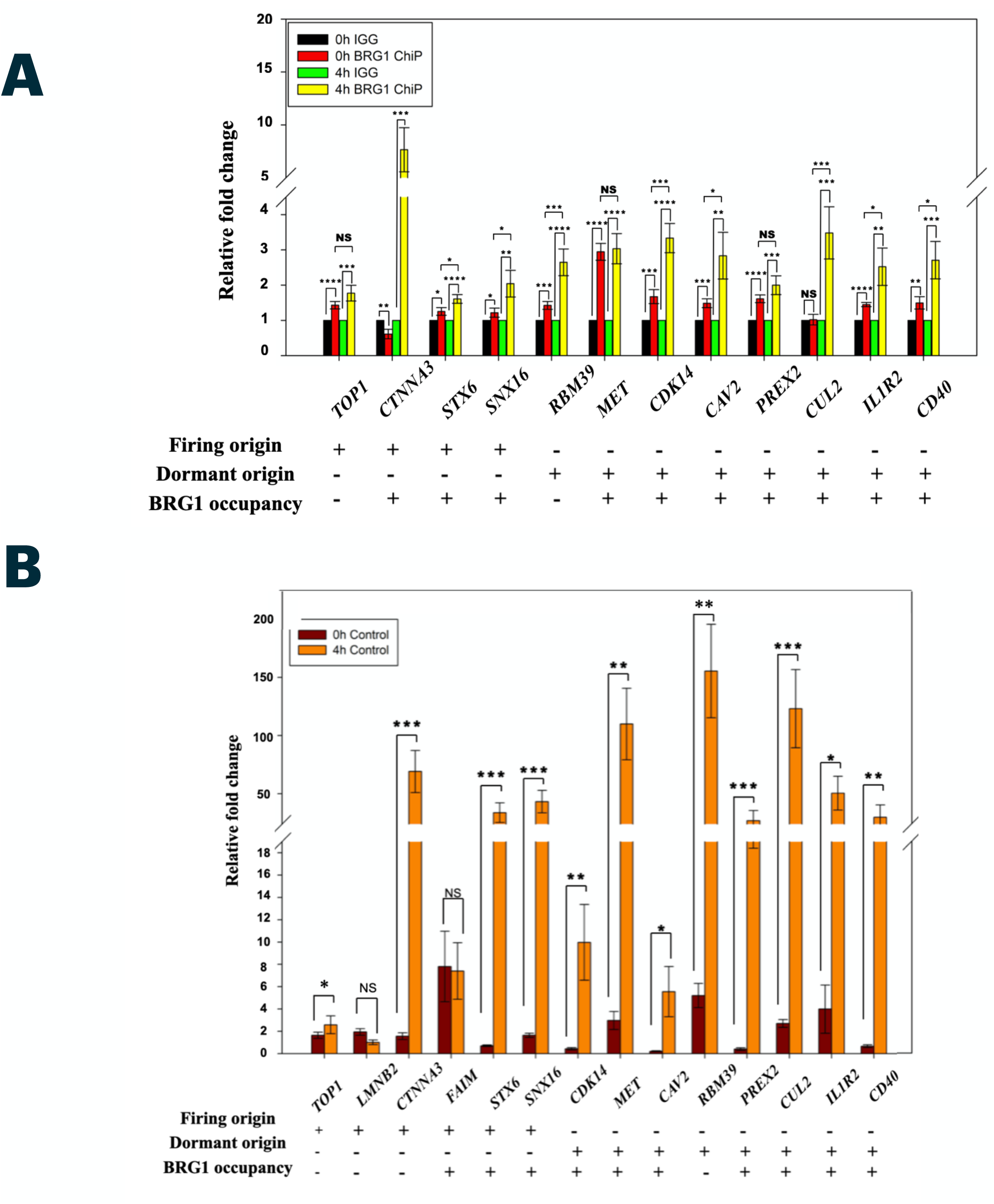
BRG1 occupancy and replication progression on selected replication origins. (A) BRG1 occupancy on selected replication origins was measured in double thymidine synchronised cells at G1/S and S phase using ChIP-qPCR. (B) CldU incorporation on selected replication origins was studied using ChIP in G1/S and S phase synchronised cells. All experiments are presented as average ± SEM of three independent biological replicates. The statistical significance (*p <0.05) was calculated using an unpaired student’s t-test.

**Figure S.4:**
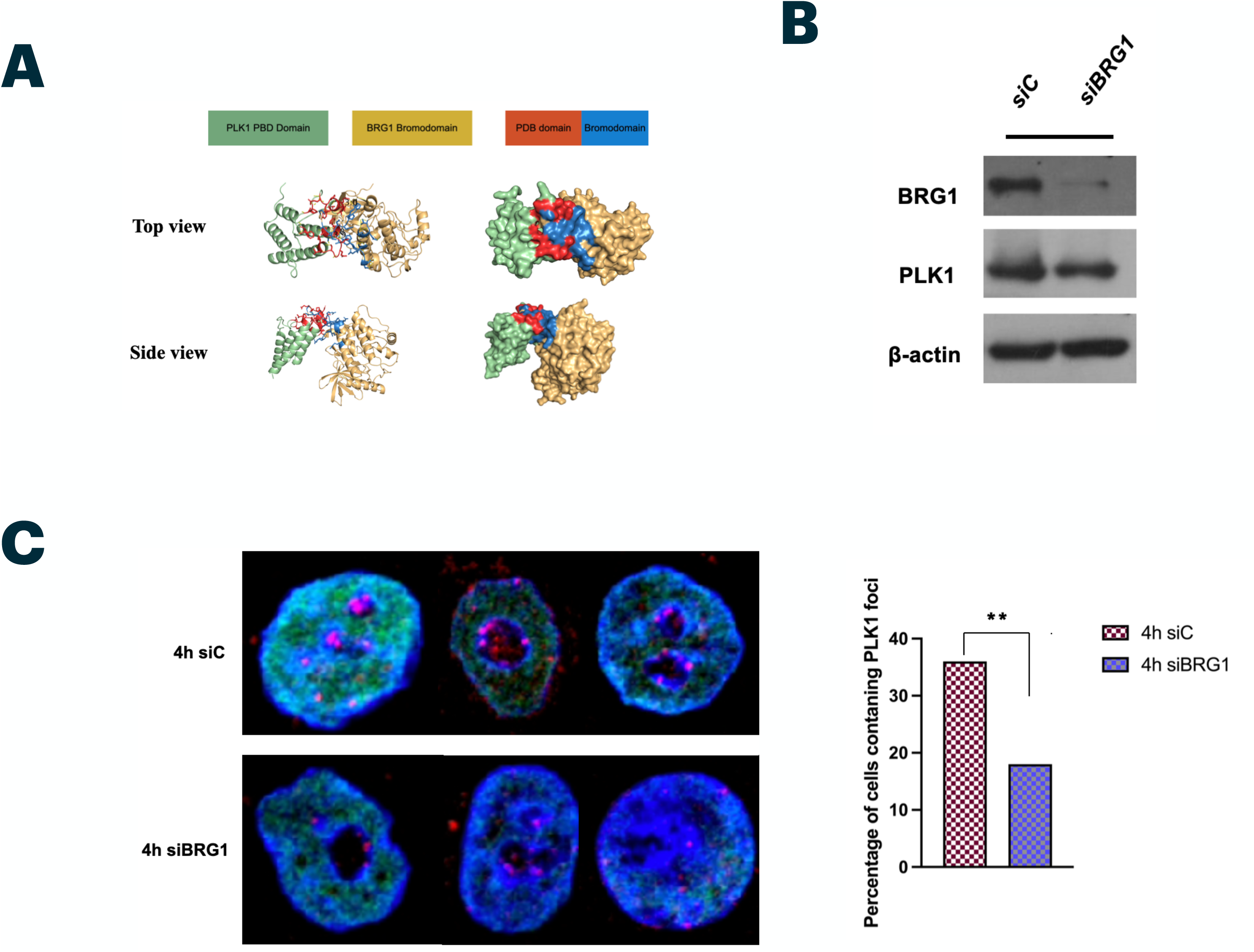
Regulation of BRG1 by PLK1 at G1/S and S phase of the cell cycle. (A) Western blot of G1/S and S phase synchronised HeLa cells transfected with vector only or wild type BRG1 and (B) Protein expression of BRG1 and PLK1 is measured in asynchronous *BRG1* depleted HeLa cells. (C) Measurement of PLK1 foci (pink) on chromatin in S phase nuclei in control (upper panel) and *BRG1* depleted (lower panel) HeLa cells. The statistical significance (*p <0.05) was calculated using an unpaired student’s t-test. For PLK1 foci quantification, (n) > 200 cells were taken from three biological replicates.

**Figure S.5:**
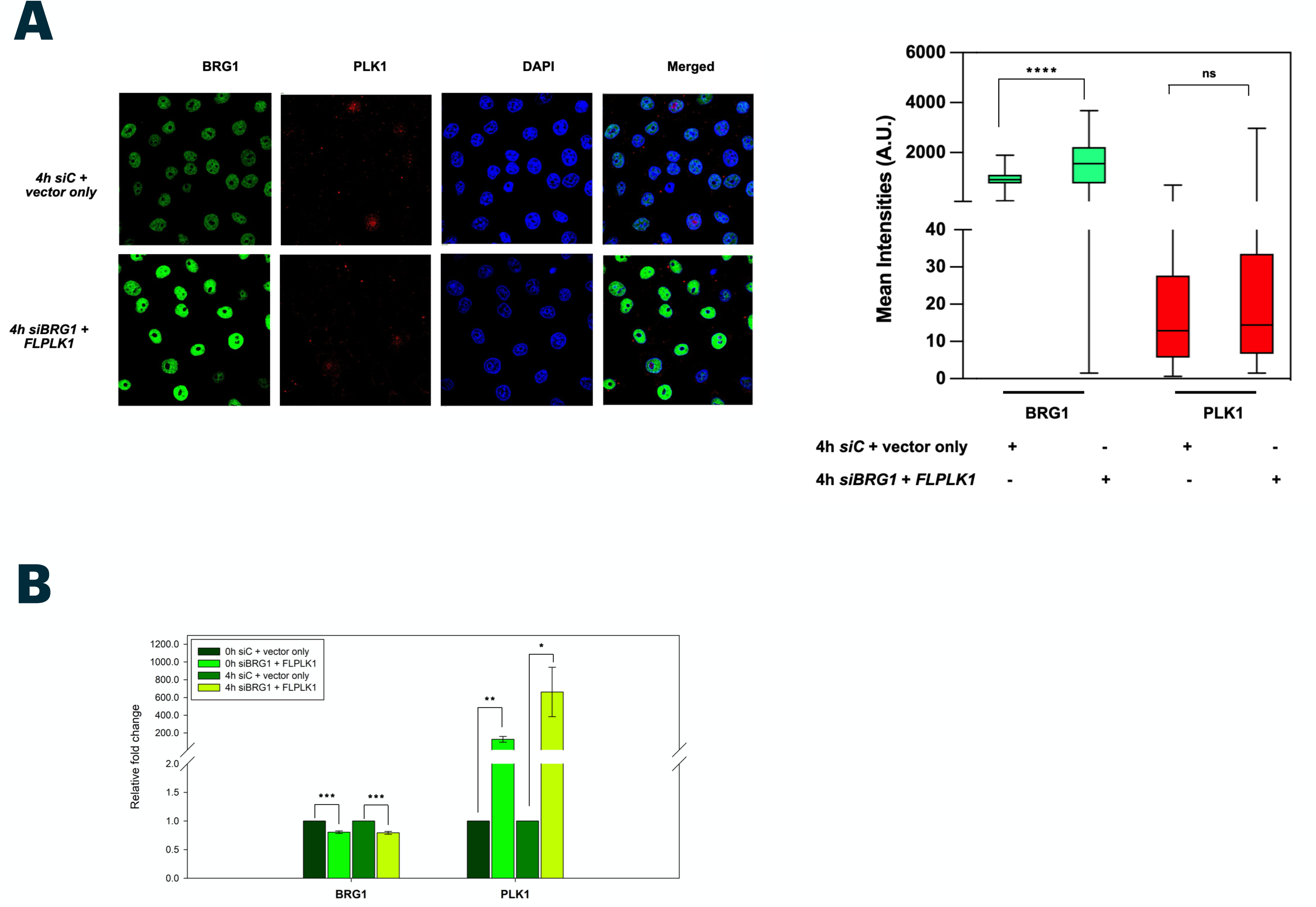
PLK1 over expression in siBRG1 cells elevates BRG1 protein. (A) Nuclear intensities of BRG1 and PLK1 protein in siC plus vector only (upper panel) and si*BRG1* plus PLK1 overexpressed S phase synchronised cells (lower panel). mRNA (B) and protein (C) expression of BRG1 and PLK1 in control (siC+vector only) and si*BRG1 PLK1* overexpressed *G1*/S and S phase synchronised cells. The box plot shows the mean distribution of intensities and statistical significance (*p <0.05) was calculated using an unpaired student’s t-test. For confocal image quantification, n > 200 cells were taken from three independent biological replicates.

**Table 1.1:**
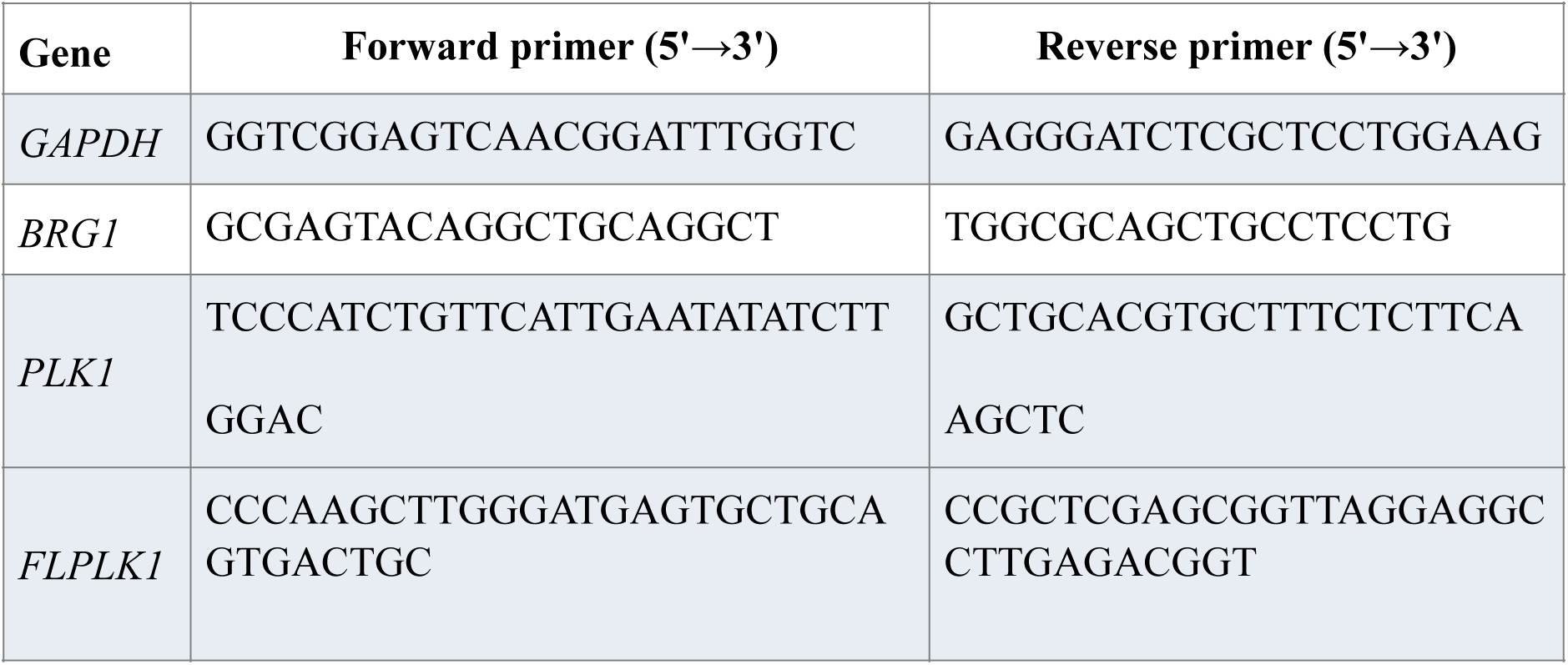
List of primers used for qRT-PCR and PLK1 cloning.

**Table 1.2:**
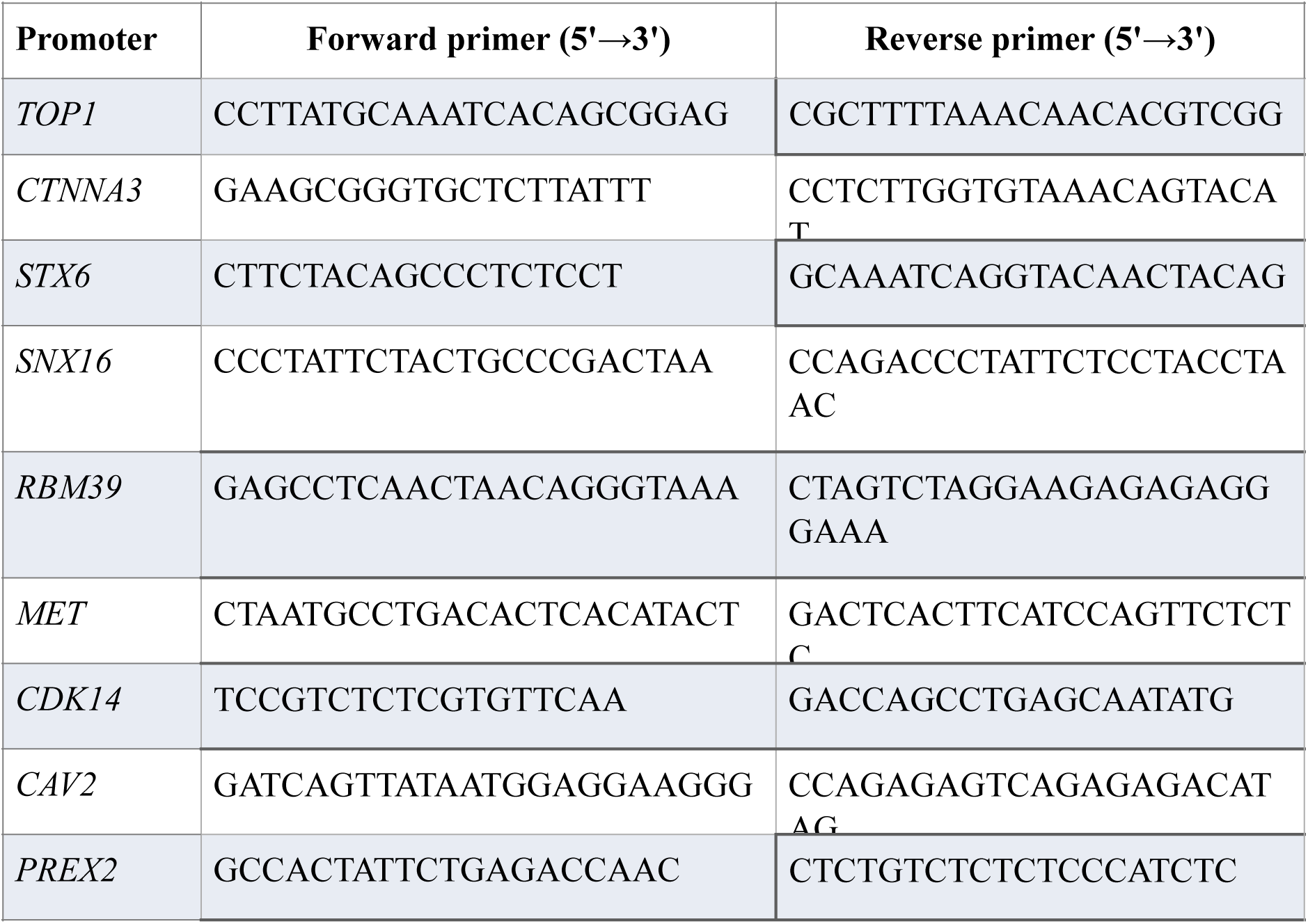
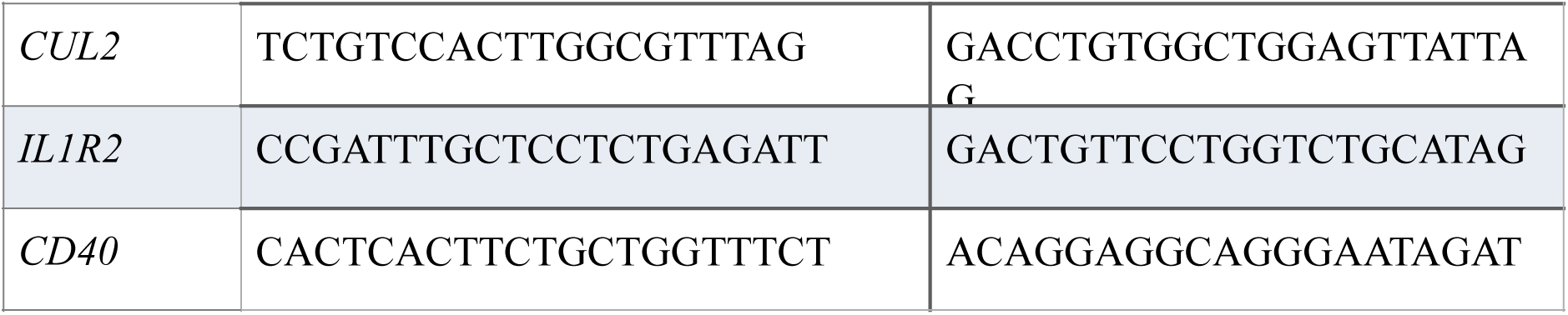
List of primers used for CldU-ChIP.

